# Adipose microenvironment promotes hypersialylation of ovarian cancer cells

**DOI:** 10.1101/2024.05.13.593990

**Authors:** Alexandra Fox, Garry D. Leonard, Nicholas Adzibolosu, Terrence Wong, Roslyn Tedja, Sapna Sharma, Radhika Gogoi, Robert Morris, Gil Mor, Charlie Fehl, Ayesha B. Alvero

**Author notes:** These authors contributed equally to this work and are listed alphabetically (Alexandra Fox, Garry D. Leonard). **Correspondence:** Ayesha B. Alvero 275 E. Hancock St. Detroit, MI 48201; Charlie Fehl 5101 Cass Avenue Detroit, MI 48202.

## Abstract

Sialylation, the addition of negatively charged sialic acid sugars to terminal ends of glycans, is upregulated in most cancers. Hypersialylation supports multiple pro-tumor mechanisms such as enhanced migration and invasion, resistance to apoptosis and immune evasion. A current gap in knowledge is the lack of understanding on how the tumor microenvironment regulates cancer cell sialylation. The adipose niche is a main component of most peritoneal cancers’ microenvironment. This includes ovarian cancer (OC), which causes most deaths from all gynecologic cancers. In this report, we demonstrate that the adipose microenvironment is a critical regulator of OC cell sialylation. *In vitro* adipose conditioning led to an increase in both ⍺2,3- and ⍺2,6-linked cell surface sialic acids in both human and mouse models of OC. Adipose-induced sialylation reprogramming was also observed *in vivo* from intra-peritoneal OC tumors seeded in the adipose-rich omentum. Mechanistically, we observed upregulation of at least three sialyltransferases, ST3GAL1, ST6GAL1 and ST3GALNAC3. Hypersialylated OC cells consistently formed intra-peritoneal tumors in both immune-competent mice and immune-compromised athymic nude mice. In contrast, hyposiaylated OC cells persistently formed tumors only in athymic nude mice demonstrating that sialylation impacts OC tumor formation in an immune dependent manner. To our knowledge, this is the first demonstration of the effect of adipose microenvironment on OC tumor sialylation. Our results set the stage for translational applications targeting sialic acid pathways in OC and other peritoneal cancers.

## 1 Introduction

Sialylation, the addition of negatively charged sialic acid sugars on terminal ends of glycans, is upregulated in most cancers and implicated across nearly all phases of cancer progression[1]. Sialic acids are 9-carbon hexosamine sugars that cap the ends of glycan chains on proteins and lipids, with roles in altered adhesion and invasion, resistance to apoptosis and immune evasion.[2; 3; 4] Sialic acids are added to growing glycan chains by twenty different sialyltransferase (STase) enzymes, which fall under one of four groups: ST3GAL and ST6GAL add sialic acid to galactose; while ST6GALNAC and ST8SIA add sialic acid to N-acetylgalactosamine or sialic acid, respectively [5]. STases add sialic acids either through an ⍺-2,3 (in the case of ST3GAL), ⍺-2,6 (in case of ST6GAL and ST6GALNAC), or ⍺-2,8 (in case ST8SIA) linkage, which refers to the stereochemistry and position of sialic acid relative to the preceding sugar residue. Upregulated expression of STases in cancers have been shown to occur through DNA hypomethylation, gene amplification or as a result of oncogene activity [5]. Hypersialylation has been correlated with pro-tumor functions in various cancer types including [6; 7; 8]. A current gap in knowledge is the lack of clear understanding on how the tumor microenvironment regulates cancer cell sialylation [9].

Peritoneal cancers such as pancreas, colon, gastric, and ovarian exhibit strong predilection to adipose rich niches in the peritoneal cavity[10; 11; 12; 13]. These sites include the adipose-rich omentum as well as the mesenteric and perigonadal adipose[14]. Adipose tissues not only serve as an energy depot that can sustain the energy requirements of rapidly growing cancer cells, but can also exert paracrine and endocrine effects by secreting adipokines, cytokines and chemokines that can support cancer cell migration and invasion[15]

Ovarian cancer (OC), by mortality rate, is the deadliest of all gynecological cancers[16; 17] A key driver of mortality is that OC is often diagnosed at a late, already metastatic stage[16; 17]. The adipose-rich omentum is an early and primary site of OC metastasis [18; 19; 20; 21]. IL-8 secreted by adipocytes has been shown to chemoattract OC cells very early in the process of metastasis formation [21]. Within the adipose niche, cross talk between adipocytes and OC cells leads to metabolic reprogramming in both cell types, which provide OC cells the required energy to sustain rapid cancer growth. Moreover, the adipose microenvironment has been shown to confer chemoresistance through Akt [22] and Bclxl pathways [23]. Following treatment, the adipose microenvironment is also a frequent site of residual and recurrent OC [24; 25; 26]. The importance of the adipose microenvironment in OC progression is underscored in studies demonstrating that the extent of tumor debulking in the adipose-rich omentum and the response of adipose-associated metastatic disease to chemotherapy is directly proportional to patient survival [24],[27]

In this study, we demonstrate that the adipose microenvironment is a critical regulator of OC cell sialylation. Using *in vitro* and *in vivo* assays and both human and mouse models of OC, we show that secreted factors from omental cultures can upregulate several STases and hence reprogram overall OC cell sialylation. Further, we demonstrate enhanced tumor establishment by hypersialylated OC cells in an immune dependent manner. Our results demonstrate that adipose-induced sialylation reprogramming has significant clinical implications in the targeting of sialylation as therapy for OC.

## 2 Materials and Methods

### 2.1 Human subjects

Human subject research was reviewed by Wayne State University IRB and found to not meet the definition of Human Participant Research and therefore exempted from IRB oversight. Samples were collected after obtaining informed consent and de-identified by the Karmanos Cancer Institute Biobanking and Correlative Sciences Core. Omentum samples were consecutively collected from patients undergoing laparoscopic or open surgery for a benign or malignant gynecological condition irrespective of diagnosis or age.

### 2.2 Cell lines and culture conditions

R182 and OCSC1-F2 human OC cell lines were established as previously described [28; 29; 30; 31; 32; 33; 34; 35; 36; 37]. A2780 (RRID:CVCL_0134) human OC cell line was a kind gift from Dr. TC Hamilton [38]. OVCAR3 (RRID:CVCL_0465) and OVCA432 (RRID:CVCL_3769) human OC cell lines were obtained from ATCC (Manassas, VA). All human cell lines were maintained in Roswell Park Memorial Institute (RMPI 1640) media containing 10% fetal bovine serum (FBS), 1% penicillin-streptomycin, 1% MEM-NEAA, 1% HEPES and 1% sodium pyruvate. Triple knock out (TKO) mouse OC cells were kindly provided by Dr. M. Matzuk [39]. TKO cells were obtained from spontaneously formed high-grade serous ovarian tumors in mice with conditional KO of Dicer and PTEN and gain of function p53 mutation (*p53*^LSL-R172H^/+*Dicer*^flox/flox^*Pten*^flox/flox^ *Amhr2*^cre/+^). TKO cells were cultured in 1:1 Dulbecco’s modified eagle medium (DMEM) and Ham’s F12 (F12) medium containing 10% fetal bovine serum and 1% penicillin-streptomycin. ID8*^Trp53-/-^* mouse OC cells (clone F3) were kindly provided by Dr. I. McNeish [40; 41] and maintained in DMEM high Glucose (Thermo Fisher Scientific, Waltham, MA) supplemented with 4% FBS, 1% Penicillin-Streptomycin, 1% Sodium Pyruvate, and 1% Insulin-Transferrin-Selenium. ID8*^Trp53-/-^* cells were derived from wild-type ID8 mouse OC cells (RRID:CVCL_IU14) by KO of p53 using CRISPR/Cas9 [40; 41]. mCherry fluorescence was stably expressed in OCSC1-F2 and TKO cells using lentivirus as previously described [35]. All cells were maintained in standard culture conditions at 37°C with 5% CO_2_. All cell lines were frequently tested for *Mycoplasma* and authenticated at least once a year by short tandem repeat (STR) profiling and used within 8 passages for each experiment.

### 2.3 Generation of human adipose conditioned media

Adipose conditioned media (ACM) were prepared as previously described [42; 43]. Briefly, 0.5 g of omentum tissue was minced with sterile razor blades and cultured in 10 mL DMEM/F12 media supplemented with 1% exosome-depleted fetal bovine serum (System Biosciences, Palo Alto, CA). ACM was collected the following day, centrifuged at 1500 RPM for 5 minutes, and stored at -80 °C until use.

### 2.4 RNA sequencing and data analysis

mRNA-seq primed from the polyA was used to determine expression profiles. Lexogen’s QuantSeq 3’mRNA-seq Library Prep Kit (FWD for Illumina) was utilized for building RNA-seq libraries from 0.1-200 ng of total RNA in 5 µl of nuclease-free ultrapure water. Libraries were quantified on the Qubit and Agilent 2200 Tapestation using the DNA High Sensitivity Screen tape. The electrophoretogram, RNA Integrity Number (RIN), and the ratio of the 28S:18S RNA bands are collectively examined to determine overall quality of the RNA. The barcoded libraries were multiplexed at equimolar concentrations and sequenced with 75 bp reads on an Illumina NovaSeq SP flow cell. Data was demultiplexed using Illumina’s CASAVA 1.8.2 software. After read quality was assessed [44], reads were aligned to the human genome (Build hg38) [45] and tabulated for each gene region [46]. Differential gene expression analysis was used to compare transcriptome changes between conditions using a paired design [47]. Significantly altered genes (p-value ≤ 0.05) were input in iPathwayGuide (Advaita Bioinformatics, Ann Arbor, MI) to identify differentially regulated Pathways.

### 2.5 Protein lysis, SDS-PAGE and Western blot analysis

Whole cell protein lysates were isolated by resuspending cell pellets in 1x Cell lysis buffer (Cell Signaling Technologies, Danvers, MA) with added Complete^TM^ Protease Inhibitor Cocktail (Millipore Sigma, Burlington, MA), followed by centrifugation for 20 minutes at 13,000 rpm. Protein lysates were quantified using BCA assay. 50 μg of protein lysate was electrophoresed on 12% SDS-polyacrylamide gels and transferred to PVDF membranes (EMD Millipore, Burlington, MA). After blocking with 5% milk, membranes were probed overnight with primary antibodies at 4°C and incubated with an appropriate secondary antibody for 1 hour at room temperature. The blots were developed using enhanced chemiluminescence and imaged using GE ImageQuant LAS 500 chemiluminescence (Cytiva Life Sciences, Marlborough, MA). The following antibodies were used: ST3GAL1 (RRID: AB_3096968) and GAPDH (RRID:AB_1078991).

### 2.6 Click chemistry

Cells were treated with 50 µM Ac_4_ManNAz for 24hrs prior to incubation with 50 µM DBCO-AF488 (Lumiprobe) for 1 hr. Cells were then rinsed with PBS containing 1% FBS prior to imaging using a Cytation 5 instrument (Agilent BioTek). Mean fluorescence was calculated using Gen5 software (RRID:SCR_017317).

### 2.7 Lectin staining and flow cytometry

Cells from culture were collected by trypsinization. Cells from tumors were dissociated using razor blades and passed through 70μm filter to obtain single cell suspension. 1×10^6^ cells were resuspended in 100 µL FACS buffer (1X PBS + 1% bovine serum albumin + 0.05% sodium azide) and stained for 30 mins on ice with the following lectins at 1:400 dilution: SNA-FITC (Vector Laboratories; RRID:AB_2336719), Mal-I-FITC (Bioworld 21761036), Mal-II-FITC (Bioworld 21511103) and PNA-FITC (RRID:AB_2315097). Pe-Cy7 conjugated anti-CD45 (RRID:AB_312979) was used at 1:100 dilution. After staining, cells were rinsed 3 times with FACS buffer. Data were acquired using CytoFLEX analyzer (RRID:SCR_019627) and CytExpert (RRID:SCR_017217) acquisition software (Beckman Coulter, Brea, CA). Data were analyzed and histograms were generated using FlowJo (RRID:SCR_008520; Becton, Dickinson and Company, Ashland, OR). For flow cytometry-assisted cell sorting (FACS) of TKO cells, SNA-stained cells were sorted using SH800S (RRID:SCR_018066; Sony Biotechnology, San Jose, CA). Cells were recovered in FBS-containing media appropriate for each cell type, washed with PBS, and plated into T25 tissue culture flasks for expansion and analysis.

### 2.8. RNA extraction and RT-qPCR

RNA was extracted using RNeasy kit (Qiagen, Germantown, MD) following manufacturer’s instructions. One μg RNA was converted to cDNA using iScript cDNA synthesis kit (Bio-Rad Laboratories, Hercules, CA) and 1:10 dilution of cDNA was used for each qPCR reaction. qPCR was performed using TaqPath™ qPCR Master Mix, CG (Thermo Fisher Scientific: A55866) with the following TaqMan primers: St3Gal1 (Thermo Fisher Scientific Assay ID: Mm00501493_m1); St6Gal1 (Thermo Fisher Scientific Assay ID: Mm00486119_m1); St6GalNac3 (Thermo Fisher Scientific Assay ID: Mm01316813_m1); and RPS17(Thermo Fisher Scientific Assay ID: Mm01314921_g1). qCPR was run on CFX96TM PCR detection system (Bio-Rad, Hercules, CA) using the following thermocycling parameters: polymerase activation at 95°C for 20 secs followed by 40 cycles of denaturation at 95°C for 15 sec and annealing/extension at 60°C for 1 min. Relative expression was calculated using the comparative ΔΔCT method. No RT samples were used as negative control. All reactions were performed in triplicates.

### 2.9 *In vivo* studies

All the described experiments using mice were approved by Wayne State University Animal Care and Use Committee (IACUC 22–03-4474) and mice were housed at Wayne State University Division of Laboratory Animal Resources. Mouse OC cells were injected intra-peritoneally (i.p.) in 7 week old female C57BL/6 mice (RRID:IMSR_JAX:000664; Jackson Laboratories, strain 000664) at 1×10^7^ or in athymic nude mice (Inotiv (Envigo) Hsd:Athymic nude-Foxn1^nu^, strain 6905F) at 5×10^6^. mCherry fluorescence was measured by live imaging under isoflurane anesthesia twice weekly using Ami HT Imaging System (Spectral Instruments, Tucson, AZ). Mice were imaged with an Excitation of 570nm, emission of 630nm. Tumor burden was quantified using mCherry region of interest (ROI) using Aura Imaging Software (Spectral Instruments). mCherry ROI area exceeding 3.4×10^8^ photons/sec (for C57BL/6) or 5×10^8^ photons/sec (for athymic nude mice) were considered above background based on imaging of non-tumor bearing mice. Animals were sacrificed when mCherry ROI area exceeded 1×10^9^ photons/second for two consecutive images or when abdominal width reached or exceeded 3.4 cm. All animals were included in the analysis and investigators were not blinded to groupings.

### 2.10 Statistical analysis

Unpaired two-tailed Student t tests, assuming Gaussian distribution, or one-way or two-way analysis of variance (ANOVA) with multiple comparisons were used for comparison between different groups. P values of 0.05 or less were considered statistically significant. Statistical analysis was performed, and all data were graphed, using GraphPad Prism v9.3.1 (San Diego, CA; RRID:SCR_002798). Data are presented as mean ± SEM.

## 3 Results

### 3.1 Adipose upregulates ovarian cancer cell sialylation

Given the significance of the adipose microenvironment in OC progression [21; 22; 42; 48; 49] we set to identify mechanisms induced by chronic exposure of OC cells to adipose secreted factors. Since adipocytes represent the primary cell type in the omentum, we obtained adipose-conditioned media (ACM) from dissociated human omentum [42; 43], treated human A2780 OC cells for 7 days with ACM, and performed transcriptomic analysis. Of the 27,162 measured genes, we observed 593 differentially expressed genes (DEGs; p<0.05; fold-changed (FC)>0.6) relative to non-ACM-treated control cells (**Fig. 1A**). Pathway Enrichment and Pathway Impact analyses showed 11 differentially regulated pathways (**Table 1** and **Fig. 1B**) and one of them was the glycosaminoglycan biosynthesis pathway (*p*=0.036; **Fig. 1B, yellow dot**). In the glycosaminoglycan biosynthesis pathway, two genes were significantly upregulated in ACM-treated cells: *B3GNT7* (*p*=0.039), which encodes β1-3- N-acetylglucosaminyltransferase and *ST3GAL1* (*p*=0.034), which encodes a STase (**Fig. 1C**). The increase in ST3GAL1, one of several STases that catalyze the addition of sialic acid to terminal ends of glycans, was validated at the protein level using two different patient omenta (**Fig. 1D, Supp. Fig. 1**). These results suggested that the adipose environment can elevate sialylation in OC cells.

**Figure 1.**
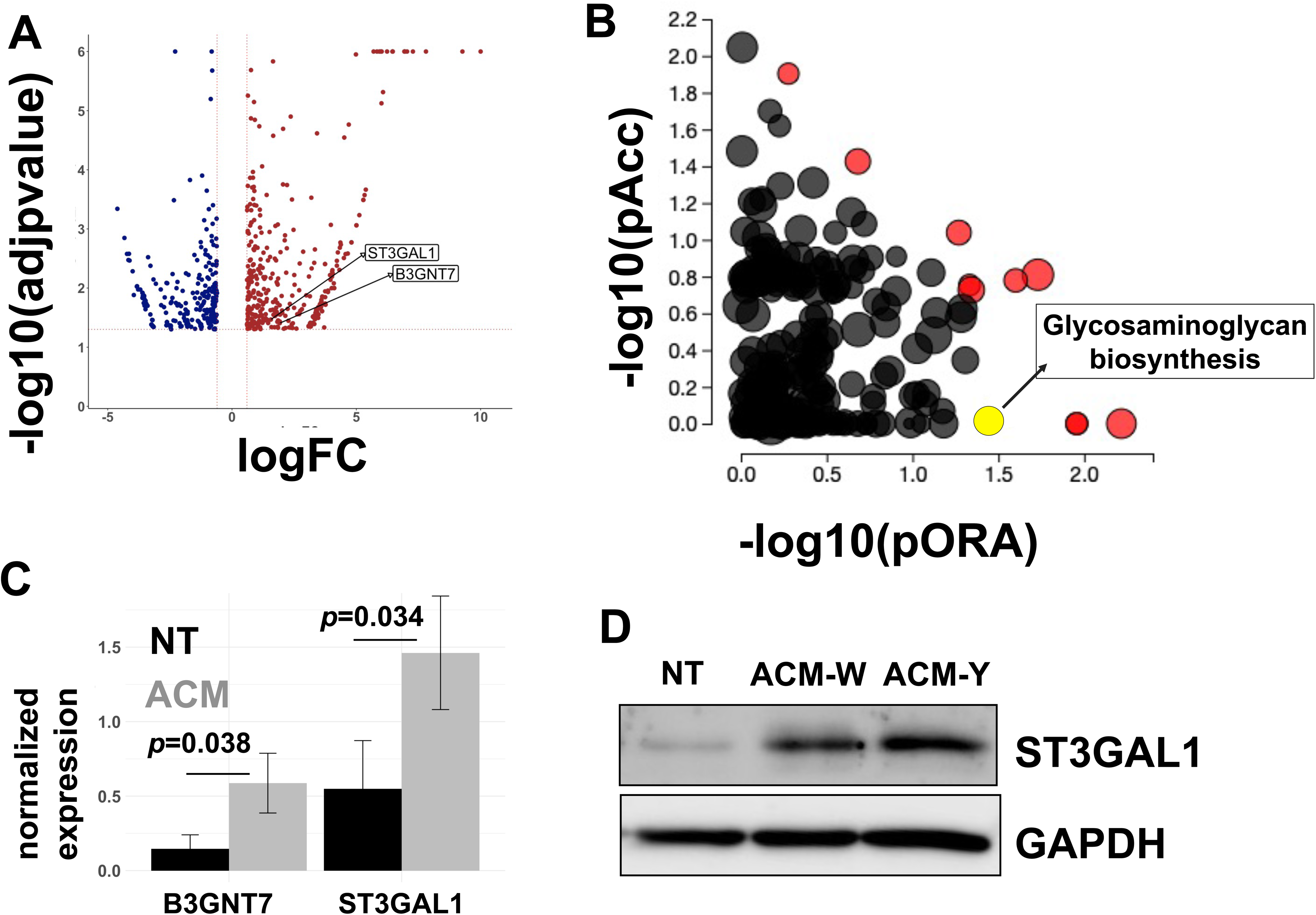
Adipose-conditioned media (ACM) upregulate sialyltransferases in human ovarian cancer cells. A2780 human OC cells were treated with ACM for 7 days prior to RNA sequencing. Control cells were maintained in growth media. **A)** Volcano plot of differentially expressed genes (DEGs; p<0.05 and fold-change>0.6) comparing Control vs ACM-treated cells; position of ST3GAL1 and B3GNT7 are shown; **B)** Differentially regulated pathways showing both Pathway impact (pORA) and Pathway enrichment (pAcc); red dots are differentially regulated and Pathway names are shown in **Table 1**; yellow dot corresponds to Glycosaminoglycan biosynthesis pathway (*p*=0.036); **C)** DEGs within Glycosaminoglycan *pathway: B3GNT7* (*p*=0.039) and *ST3GAL1* (*p*=0.034); **D)** Western blot analysis of human R182 ovarian cancer cells treated with ACM from either patient W or patient Y showing upregulation of ST3GAL1. *NT*, no treatment control.

**Table 1.** Differentially regulated pathways in ovarian cancer cells treated with adipose-conditioned media.

| Pathway name | p value |
| --- | --- |
| Ribosome | 0.006 |
| Glycine, serine and threonine metabolism | 0.011 |
| Nicotine addiction | 0.011 |
| cAMP signaling pathway | 0.019 |
| Amyotrophic lateral sclerosis | 0.027 |
| Central carbon metabolism in cancer | 0.031 |
| Glycosaminoglycan biosynthesis | 0.036 |
| SNARE interactions in vesicular transport | 0.039 |
| Melanoma | 0.045 |
| Thyroid cancer | 0.047 |
| PD-1 checkpoint pathway in cancer | 0.049 |

To determine if adipose-induced upregulation of STases leads to a measurable increase in cell surface sialylation, we used copper-independent and strain-promoted azide-alkyne click chemistry (SPAAC) to directly label sialic acid sugars. SPAAC reactions connect an azide species and alkyne via a bioorthogonal cycloaddition reaction [50; 51; 52]. The azide component was installed on sialic sugars through feeding tetraacetylated N-azidoacetyl-mannosamine (ManNAz), which is metabolized and converted to sialic acid azide in cells and added to terminal ends of sialoglycans. The strained alkyne component we used was dibenzocyclooctyne (DBCO) coupled to AF488 fluorophore (DBCO-AF488).

When DBCO-AF488 was added to cells that had metabolically incorporated azides on cell surface sialic acids, the reaction resulted in a covalent bond that linked the fluorophore to cell surface sialic acids (**Fig. 2A**). We first determined basal sialic acid expression. Click chemistry performed on mCherry+ TKO mouse OC cells showed membranal green staining demonstrating cell surface sialylation (**Fig. 2B**). To quantitate the effect of adipose conditioning on sialic acid expression, we treated R182 human OC cells with ACM for 72 h and ManNAz was added during the last 24 h of treatment (**Fig. 2Ci**). At the end of the treatment, cells were incubated with DBCO-AF488. Quantification of fluorophore signal showed significant upregulation of cell surface sialic acid expression with ACM compared to ManNaz only control (**Fig.2Cii**). These results showed that OC cells expressed basal cell surface sialoglycans, which were significantly enhanced by adipose.

**Figure 2.**
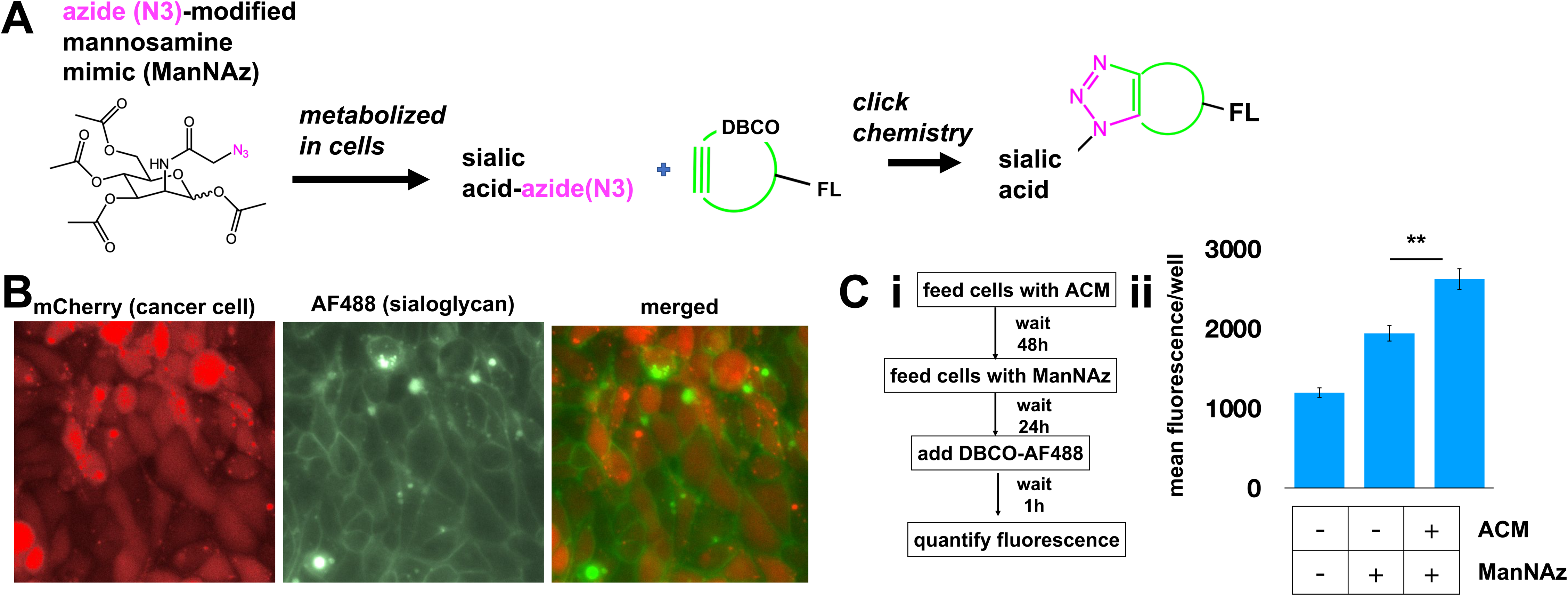
Adipose-conditioned media (ACM) upregulate cell surface sialylation in human and mouse ovarian cancer cells. **A)** Diagram of click chemistry detailed in text**; B)** mCherry+ TKO mouse OC cells were treated with 50 μM ManNAz every day for 3 days followed by treatment with DBCO-FITC. Microscopy analysis shows basal expression of cell surface sialoglycans; **C) *i,*** treatment protocol with ACM prior to click chemistry; ***ii***, R182 human OC cells were treated as in Ci and mean intensity of AF488 was quantified. Note basal sialylation, which is upregulated by ACM treatment. Data are presented as mean ± SEM (n=3); ** *p* = 0.0062 by One-Way ANOVA with *post-hoc* multiple comparison analysis.

### 3.2 Adipose upregulates α-2,6- and α-2,3-linked sialic acids on ovarian cancer cells

For a more comprehensive characterization of cell surface sialylation, we used a panel of four fluorophore-tagged lectins: *Sambucus nigra* lectin (SNA), *Maackia amurensis* Lectin I (MAL I), *Maackia amurensis* Lectin II (MAL II), and peanut agglutinin (PNA). Lectins are sugar binding proteins isolated from plants and animals and are classically used to evaluate glycan structures. SNA preferentially binds α-2,6-linked sialic acids and is a good indicator of ST6GAL1 activity [58]. MAL -I and Mal-II preferentially binds α-2,3-linked sialic acids, the enzymatic products of ST3GAL1[53; 54; 55]. Finally, PNA detects non-sialylated galactose, which is one of the required precursors to sialic acid modification on cell surface glycans [56]. We first characterized basal cell surface sialylation and used these lectins to stain a panel of human (OCSC1-F2, R182, OVCAR3 and OVCA432; **Supp. Fig. 2**) and mouse (TKO and ID8p53KO; **Supp. Fig. 3**) OC cell lines. All human cell lines showed positive staining for SNA and PNA, although with varying intensity (**Supp. Fig. 2**). Human cell lines with higher staining for SNA (i.e. R182 > OCSC1-F2) showed lower staining for PNA, as expected. MAL-I staining was only observed in OVCAR3 and MAL-II staining was observed in R182, OVCAR3 and OVCA432 but not in OCSC1-F2. Sialylation pattern on the two mouse cell lines tested was also variable. Both mouse cell lines showed positive staining for SNA and PNA (**Supp. Fig. 3**). Only TKO showed positive staining for MAL-II. Neither of the mouse cell lines stained positively for MAL-I.

Having characterized basal cell surface sialylation, we then utilized the OCSC1-F2 human OC cells to determine the effect of adipose on specific sialic acid linkages. Thus, OCSC1-F2 cells were treated for 7 days with ACM obtained from three different patient omenta prior to lectin staining. Compared to control cultures, OCSC1-F2 cells treated with ACM showed a trend of increased cell surface expression of α-2,6- and α-2,3-linked sialic acids, as detected by SNA and MAL-I staining, respectively. Despite this trend, statistical significance was not reached (**Fig. 3A,B**), suggesting that the effects had relatively high variability. Minimal increase in MAL-II staining and a decrease in PNA staining were observed in ACM-treated cells but the difference from control cells was also not statistically significant (**Fig. 3C,D**).

**Figure 3.**
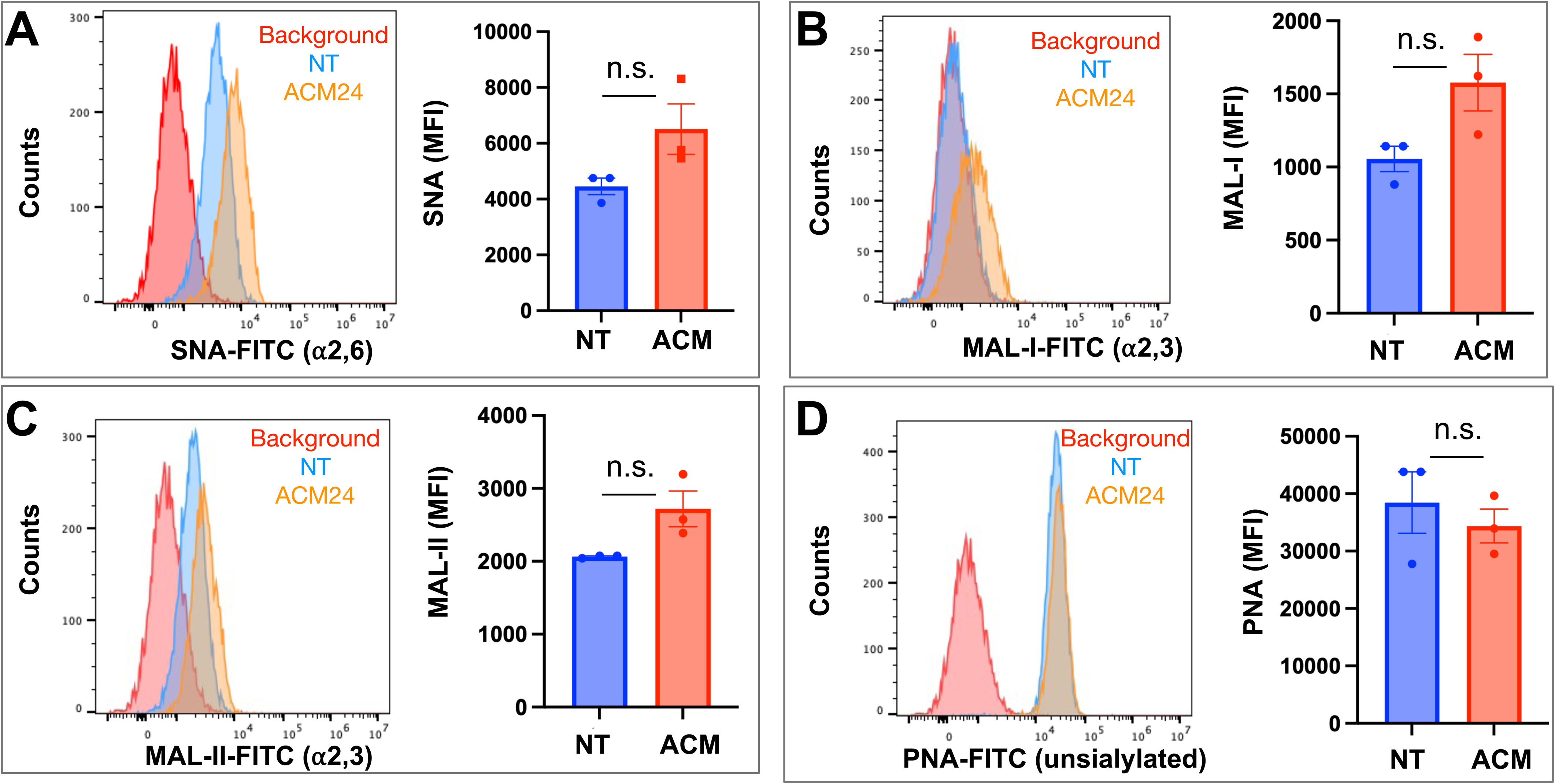
Adipose-conditioned media (ACM) upregulate ⍺2,6 and ⍺2,3 sialic acids in human ovarian cancer cells. OCSC1-F2 human OC cells were treated with ACM from three different patients (ACM 22, ACM24, ACM 26) for 7 days prior to staining with **A)** FITC-tagged SNA; **B)** FITC-tagged MAL-I; **C)** FITC-tagged MAL-II; **D)** FITC-tagged PNA. Histograms show results from ACM24. Graphs show mean ± SEM.

We noted in our lectin staining panels that TKO mouse OC cells reproducibly generated two distinct SNA-staining populations *in vitro* (**Supp. Fig. 3**). Over time, the percentage of cells in these two sub-populations fluctuated between ca. 20-60%, but the two distinct populations were persistent and were maintained in culture when followed until 9 passages (**Supp. Fig. 4**). Flow cytometry assisted cell sorting (FACS) allowed us to further interrogate sialylation on these two cell subpopulations. After authentication through STR profiling (**Supp. Fig. 5**), lectin staining comparing TKO^SNAhigh^ and TKO^SNAlow^ cells showed that these cultures were only different in SNA staining for α-2,6-linked sialic acids. They demonstrated comparable staining for α-2,3-linked sialic acids via MAL-I and MAL-II. TKO^SNAhigh^ cells showed slightly lower PNA staining compared to TKO^SNAlow^ cells (**Supp. Fig. 6**). To further demonstrate the effect of adipose secreted factors, we treated TKO^SNAlow^ cells with ACM. We noted an increase in SNA and MAL-I in ACM-treated TKO^SNAlow^ cells (**Fig. 4**), which parallels what was observed with ACM-treated OCSC1-F2 human OC cells (**Fig. 3**). In TKO^SNAlow^ cells, we also noted a decrease in PNA staining upon treatment with ACM (**Fig. 4**). No changes were observed with MAL-II staining. Taken together, these results demonstrate that adipose secreted factors can upregulate both α-2,6- and α-2,3-linked sialic acids in both human and mouse ovarian cancer cells.

**Figure 4.**
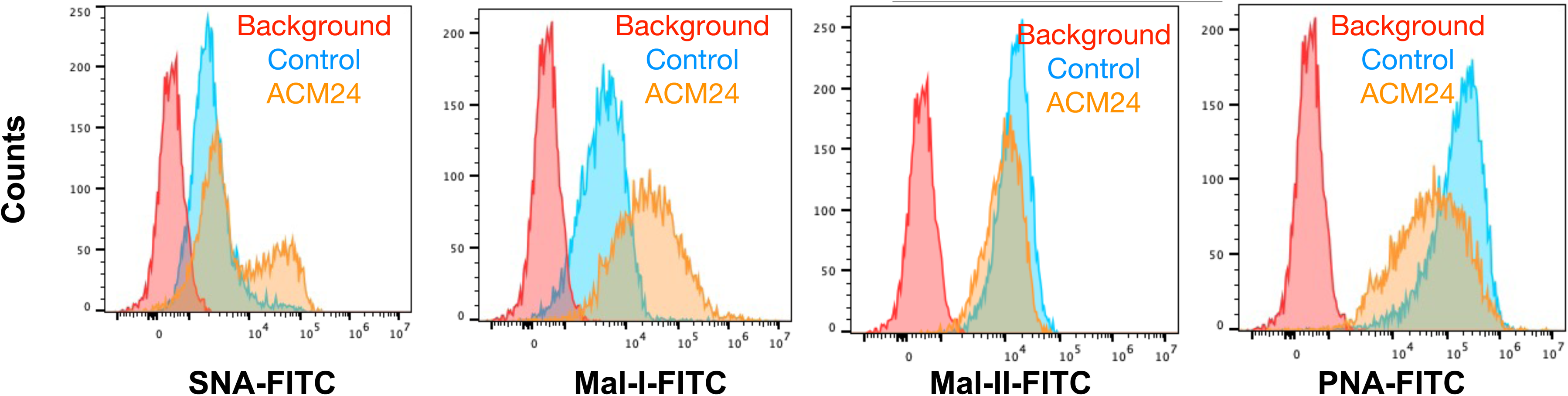
Adipose-conditioned media (ACM) upregulate ⍺2,6 and ⍺2,3 sialic acids in mouse OC cells. TKO^SNAlow^ cells were sorted by FACS and treated with ACM24 for 7 days prior to lectin staining.

### 3.3 *In vivo* engraftment reprograms ovarian cancer cell sialylation

To determine if the adipose-induced increase in sialylation observed *in vitro* is recapitulated *in vivo*, we established i.p tumors from parental mCherry+ TKO mouse OC cells in C57BL/6 mice. We previously reported the characterization of i.p. ovarian tumors formed by this model and showed its preferential seeding to omentum, pelvic fat, and mesenteric adipose [57; 58]. Necropsy showed omental implants (**Fig. 5**) as previously reported [57; 58]. We then compared cell surface sialic acid expression between TKO cancer cells in culture (**Fig. 5, top panel**) and dissociated TKO cancer cells from the omentum implants (**Fig. 5, bottom panel**). Interestingly, we observed sialylation reprogramming upon *in vivo* engraftment. Unlike TKO cells in culture, which showed two peaks for SNA staining, TKO cells from dissociated omental tumors showed a single SNA peak, which matched the staining intensity observed in the TKO^SNAhigh^ cell population (**Fig. 5**). In addition, TKO cells from dissociated tumors showed increase in both MAL-I and MAL-II staining and decrease in PNA compared to TKO cells in culture. These results demonstrate that *in vivo* engraftment may favor or select TKO^SNAhigh^ cells. Additionally, these data demonstrate a broad reprogramming of sialylation in ovarian tumors with increase in both α-2,6- and α-2,3- sialic acid linkages.

**Figure 5.**
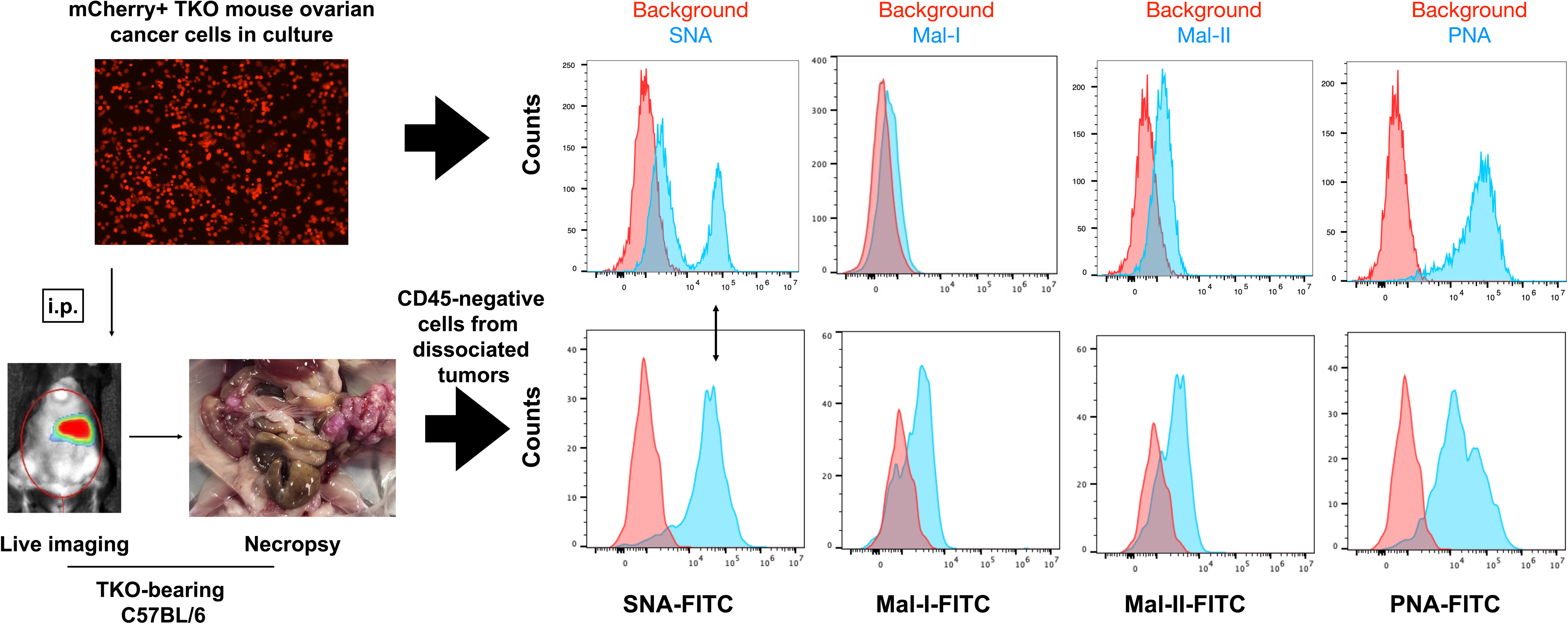
*In vivo* engraftment upregulates general sialylation. *top panel*, Cell surface sialic acid expression in mCherry+ TKO mouse OC cells in culture as detected by SNA, MAL-I, MAL-II and PNA; *bottom panel*, mCherry+ TKO mouse OC cells were injected i.p. in C57BL/6 mice and omental tumors were dissociated, stained with anti-CD45 and FITC-tagged lectins (n=5). Histograms show FITC staining from CD45-negative population. Note loss of SNA-low population and increase in Mal-I and Mal-II upon *in vivo* tumor formation. Histograms show data from one mouse. Similar results were observed in other mice. Double sided arrow shows SNA levels in tumors is comparable to SNA levels in TKO^SNAhigh^ cells.

### 3.4 Heterogeneous pool of sialylated ovarian cancer cells in culture

The finding in TKO mouse OC cultures of two subpopulations of cells based on SNA staining is in line with previous reports of heterogeneity in expression of α-2,6 sialic acids in breast and lung cancer cultures [59; 60; 61]. We further confirmed that these cells have differential surface sialic acid levels by treating TKO^SNAhigh^ and TKO^SNAlow^ cells with neuraminidase. Neuraminidase removes all cell surface sialic acids and exposes the underlying galactose, which can be bound by PNA (**Supp. Fig. 7A**). Indeed, baseline PNA staining showed that TKO^SNAhigh^ cells had lower PNA staining compared to TKO^SNAlow^ cells (**Supp. Fig. 7B**) and thus suggests higher cell surface sialic acid in TKO^SNAhigh^ cells. After neuraminidase treatment however, both cell populations showed comparable PNA staining further proving initial difference in cell surface sialic acid levels between the two cell subpopulations.

We then further characterized these two cell subpopulations (**Fig. 6A**) and performed RNA sequencing to identify key genes that may regulate the sialylation differences. We focused on the expression of 30 sialylation-associated genes [62] and found significant difference in expression of *St3gal1* (FDR=0.0003), *St3gal5* (FDR=5×10^-5^), *St6gal1* (FDR=1×10^-125^), *St6galnac3* (FDR=0.0007), *St8sia1* (FDR=0.042), and *Slc35a1* (FDR=0.004). Of these DEGs, *St6gal1* and *St6galnac3* demonstrated the highest fold increase in TKO^SNAhigh^ compared to TKO^SNAlow^ cells of up to 14-fold and 10-fold increase, respectively (**Fig. 6B**). The upregulation in both *St6gal1* and *St6galnac3* was further validated by RT-qPCR (**Fig. 6C,D**). Taken together, these data are consistent with a heterogeneous pool of sialylated OC cells in culture.

**Figure 6.**
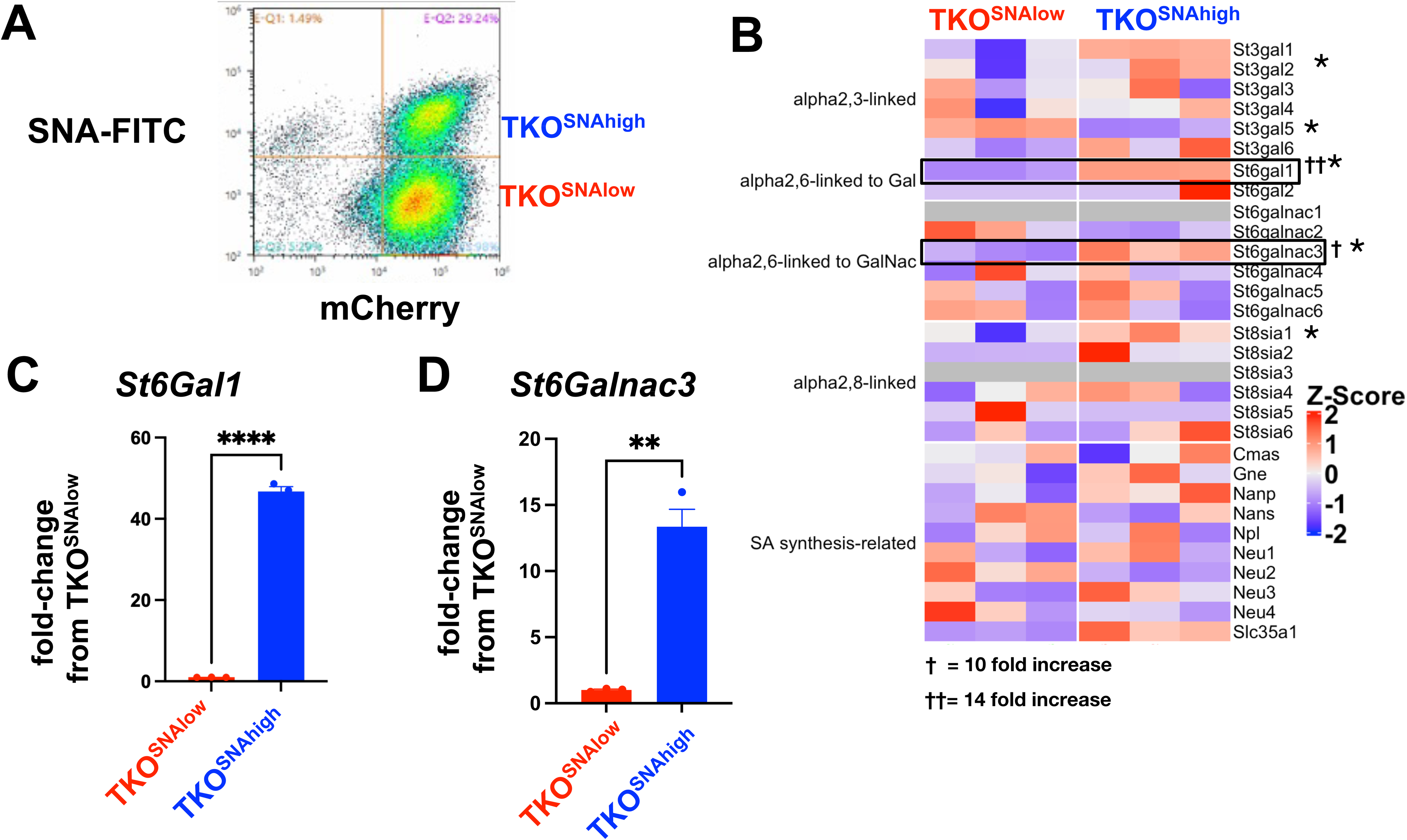
Heterogeneity of sialylation in TKO mouse OC cultures. **A)** Gating strategy for FACS to isolate TKO^SNAhigh^ and TKO^SNAlow^ cells from parental TKO cultures; **B)** Heatmap of 30 sialylation-related genes from RNA sequencing performed on TKO^SNAhigh^ and TKO^SNAlow^ cells. * denotes genes that are statistically significant (FDR<0.05). Note upregulation of *St6Gal1* (FDR=5×10^-5^) and *St6GalNac3* (FDR=0.0007). Increase in *St6Gal1* **(C)** and *St6GalNac3* **(D)** mRNA was validated by RT-qPCR. Data are presented as mean ± SEM (n=3); *** *p*<0.001, **** *p*<0.0001.

### 3.5 Hyposialylated ovarian cancer cells fail to form tumors in an immune-dependent manner

The observation that parental TKO OC cells formed i.p. tumors that consisted of only TKO^SNAhigh^ cells (**Fig. 5**) suggested that TKO^SNAlow^ cells are not tumorigenic. To test this hypothesis, we injected each subpopulation i.p. in immune-competent C57BL/6 mice. We observed tumor formation only in mice administered TKO^SNAhigh^ cells. Logarithmic tumor growth was seen in these mice beginning at day 30 (**Fig. 7A,B**). In contrast, mice injected with TKO^SNAlow^ cells demonstrated measurable disease only immediately after injection (day 3), after which point the signal dropped and all but one mouse remained disease-free until day 70 (**Fig. 7A,B**). As such, tumor growth rate was significantly different in TKO^SNAhigh^ group compared to TKO^SNAlow^ group (p<0.0001; **Fig. 7A**) and mice in TKO^SNAhigh^ group showed significantly shorter overall survival (p=0.035; **Fig. 7C**). These results show that SNA/sialic acid enriches for OC cells that are tumorigenic in immune-competent mice.

**Figure 7.**
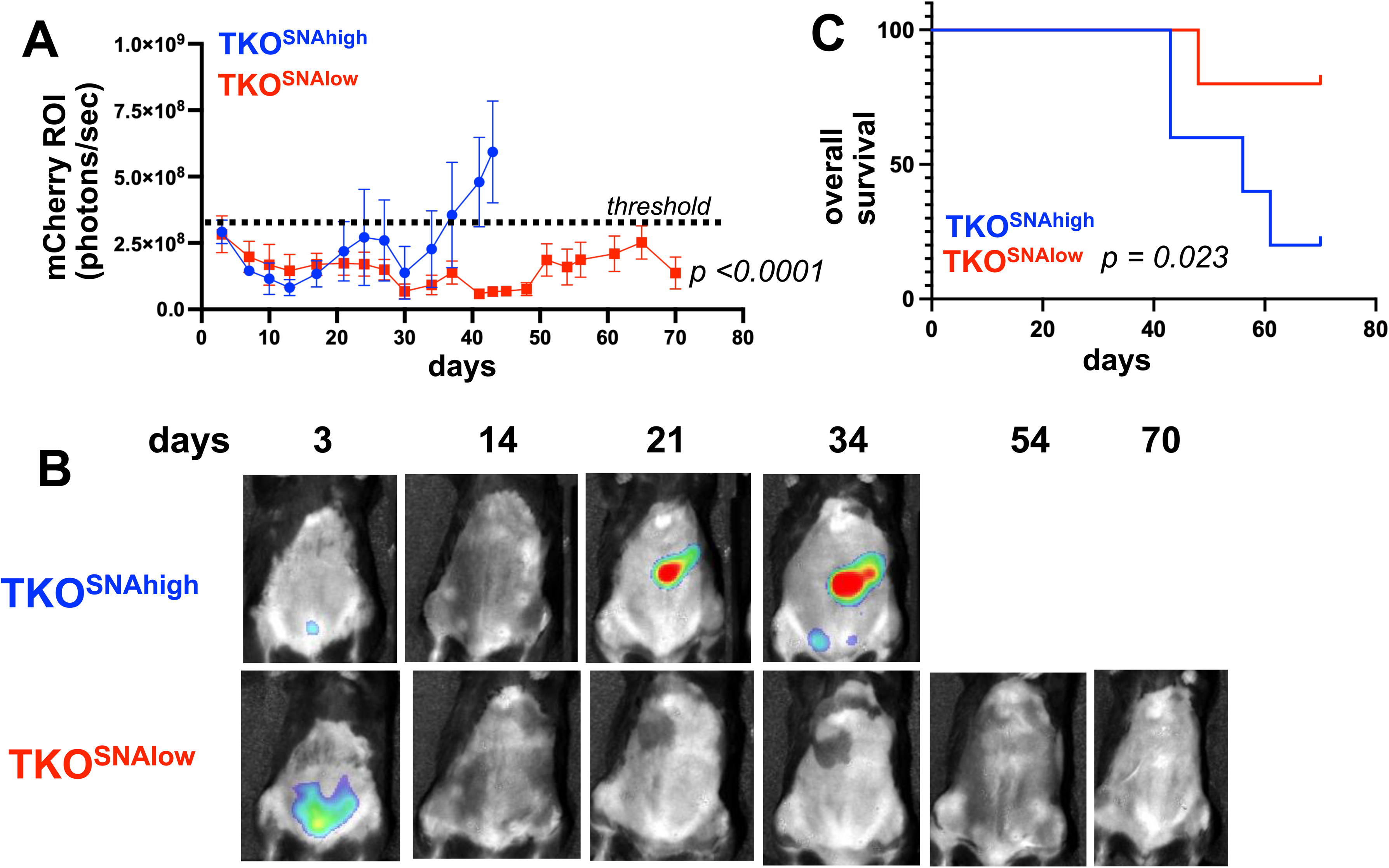
TKO^SNAlow^ cells do not form tumors in immune-competent mice. 1×10^7^ TKO^SNAhigh^ or TKO^SNAlow^ cells were injected i.p. in female C57BL/6 mice (n=5). mCherry fluorescence was acquired every 3-4 days and mCherry ROI area was quantified as measure of i.p. tumor burden. **A)** Tumor growth curves showing significant difference in measured mCherry ROI between groups (*p<0.0001* by Two-Way ANOVA). Dashed line shows threshold for mCherry signal; **B)** Representative images obtained from live imaging; **C**) Kaplan-Meir survival curve showing significantly shorter overall survival in mice injected with TKO^SNAhigh^ cells (*p*=0.023).

The observed difference in tumorigenic potential between TKO^SNAhigh^ and TKO^SNAlow^ cells in immune competent mice may be due to cell-intrinsic mechanisms (i.e. anoikis resistance, invasion capacity, metabolic fitness, etc) or these differences may be immune related. To determine the contribution of the immune system we injected each cell population in athymic nude mice lacking T cells. Interestingly, in the absence of T cells, TKO^SNAlow^ cells were able to form tumors, albeit with slower kinetics (**Fig. 8A,B**). Mice injected TKO^SNAlow^ cells showed significant delay in tumor formation (*p*=0.04) but there was no significant difference in overall survival (**Fig. 8A,C**). Taken together, our results demonstrate that the tumorigenic capacity of hyposialylated OC cells can be fully inhibited by the adaptive immune system. In contrast, the innate immune system can only delay but not fully prevent its tumorigenic potential.

**Figure 8.**
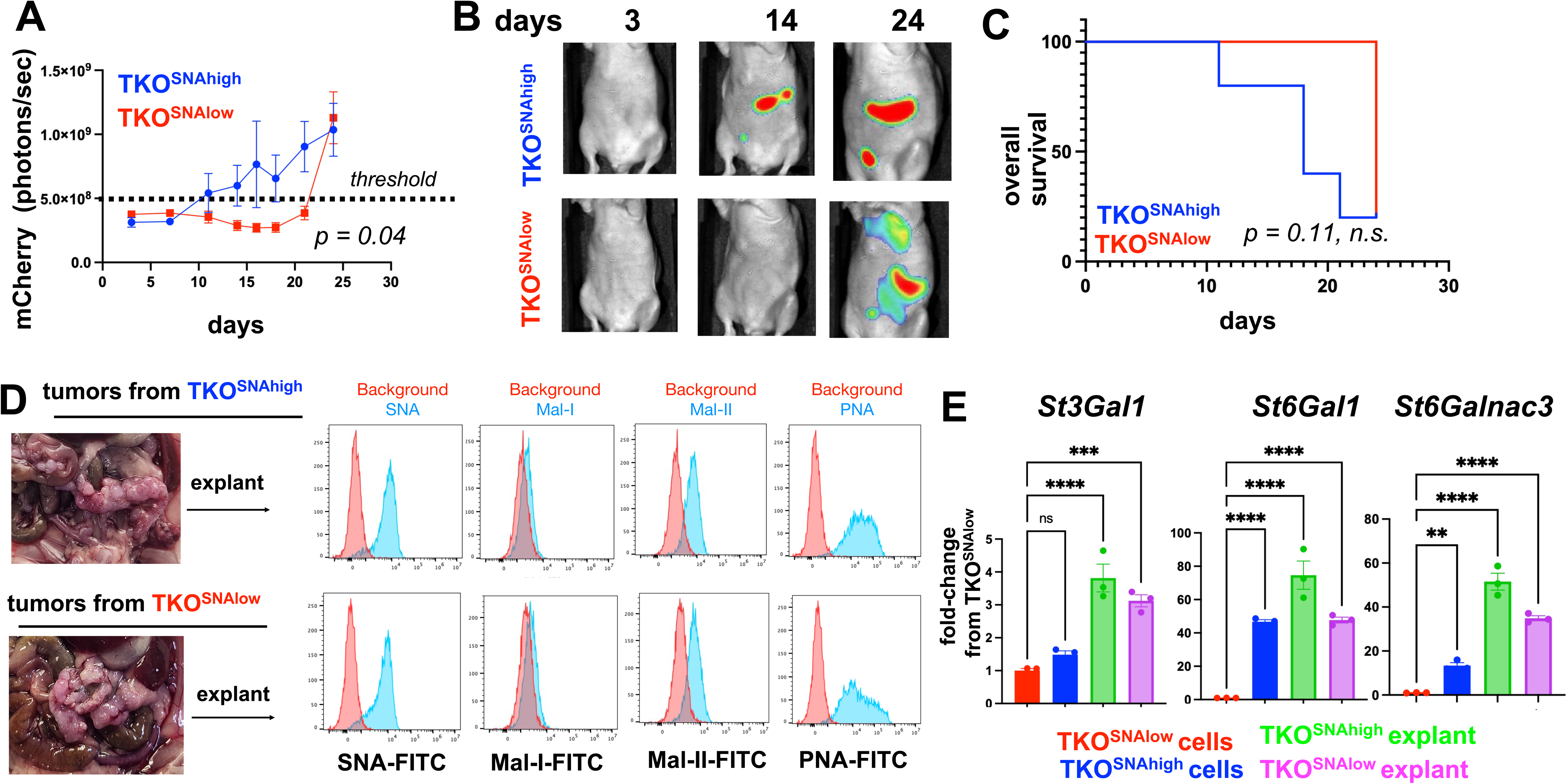
TKO^SNAlow^ cells undergo sialylation reprogramming and form tumors in immune-compromised mice. 5×10^6^ TKO^SNAhigh^ or TKO^SNAlow^ cells were injected i.p. in female athymic nude mice (n=5). mCherry fluorescence was acquired every 3-4 days and mCherry ROI area was quantified as measure of i.p. tumor burden. **A)** Tumor growth curves showing significant difference in measured mCherry ROI between groups (*p=0.04* by Two-Way ANOVA). Dashed line shows threshold for mCherry signal; **B)** Representative images obtained from live imaging; **C**) Kaplan-Meir survival curve showing no significant difference in overall survival between groups (*p*=0.11, ns); **D)** Necropsy shows omentum as primary location of i.p. tumors. Omental tumors were dissociated and cultured as explants for 2 passages prior to lectin staining; **E)** RT-qPCR for *St3Gal1*, *St6Gal1*, and *St6GalNac3.* Data are presented as mean ± SEM (n=3); ** *p* <0.01, *** *p* < 0.001, \*\*\*\**p*<0.0001 by One-Way ANOVA.

### 3.6 Successful tumor formation by hyposialylated ovarian cancer cells in immune-compromised mice leads to hypersialylation

The observation that TKO^SNAlow^ cells form tumors in immune deficient mice provided a platform to further validate *in vivo* sialylation reprogramming. Thus, we characterized the tumors formed by both the TKO^SNAhigh^ and TKO^SNAlow^ cells in athymic nude mice. Necropsy showed that majority of the i.p. tumors were seeded in the omentum (**Fig. 8D**). We then established cultures from dissociated omental explants and characterized their sialylation levels. SNA staining showed comparably high SNA intensity between explants from TKO^SNAhigh^ and TKO^SNAlow^ tumors demonstrating that TKO^SNAlow^ cells are re-programmed *in vivo* to gain α-2,6-sialylation (**Fig. 8D**). In addition to equivalent SNA staining, explants from both groups showed positive staining for Mal-II demonstrating gain in α-2,3-sialylation as well (**Fig. 8D**). Finally, we measured the levels of *St3Gal1*, *St6Gal1,* and *St6GalNac3* in the tumor explants. qPCR data showed significant increase in all STases in the tumor explants compared to TKO^SNAlow^ cells grown in culture (**Fig. 8E**). Taken together, our results demonstrate that sialylation impacts tumorigenic potential of OC cells in an immune-dependent manner and that the adipose microenvironment is an important inducer of OC cell sialylation reprogramming.

## 4 Discussion

We demonstrate in this study that the adipose microenvironment is a critical regulator of OC cell sialylation. We first took a broad approach to how secreted factors from adipose-rich omentum impacted the OC cell transcriptome. We discovered a significant effect on the STase, ST3GAL1, and using human and mouse models of OC further characterized adipose-induced sialylation. Upon adipose conditioning *in vitro*, both human and mouse OC cells increased sialylation for both α-2,3 and α-2,6 linked sialic acids. Further, *in vivo* engraftment, which for OC typically occurs in the omentum, also lead to increased overall sialylation. Mechanistically, we observed increased expression of not only St3gal1, but also St6gal1 and St6galnac3 upon *in vivo* engraftment. Intriguingly, we discovered two distinct subpopulations of OC cells with low or high α-2,6-sialic acid levels. When separately injected into immune-competent mice, these subpopulations had altered tumorigenicity and only the hypersialylated, but not the hyposialylated, OC cells exhibited consistent tumorigenic potential. Interestingly, the lack of T cells in athymic nude mice allowed tumor formation of hyposialylated OC cells. Nevertheless, upon successful tumor formation, both hypersialylated and hyposiaylated OC cells displayed increased overall sialylation.

Like most solid tumors in the peritoneal cavity, OC preferentially metastasizes to the adipose-rich omentum [63]. Upon establishment in this niche, studies have shown that OC cells undergo metabolic reprogramming characterized by upregulation of fatty acid intake, shift to β-oxidation, and diversion of glucose towards glycerol-3-phosphate [19; 21; 64; 65]. To our knowledge however, this is the first time that the effect of adipose on cancer cell sialylation has been reported. Sialic acid patterns on cell surfaces are regulated stochastically by the levels of individual STase[66; 67] There are 20 known STases, many of which are conserved in mice and humans [68]. Previous reports have shown that high-grade serous OC with high expression of sialic acid-related genes demonstrate worse overall survival [62]. Further, ST3GAL3 was identified as most predictive of prognosis [62]. Our data showed that St6gal1 and St6galnac3 are upregulated in the highly sialylated mouse OC cell subpopulations and in addition, that exposure to factors secreted by adipose tissue upregulates at least three STases, St3gal1,St6gal1 and St6galnac3. The exact mechanism by which adipose secreted factors upregulate STases and thereby increase overall sialylation is under investigation in our lab. Given the known metabolic reprogramming that occurs in OC cells in the adipose niche, we speculate that the changes in nutrient flux and metabolic signaling, which are also major determinants of glycosylation patterns in cells [69; 70], could play a role.

Cell surface sialic acids are ligands for Siglec receptors on immune cells. Siglecs have differential expression on various types of immune cells and most are immunosuppressive via a cytosolic immunoreceptor tyrosine-based inhibitor motif (ITIM) domain[71]. Sialic-acid/Siglec binding has been shown to lead to failed maturation of macrophages, generation of myeloid-derived suppressor cells (MDSC), promotion of T regulatory cells, and inactivation of natural killer cells [62; 72; 73]. There are 15 Siglecs in humans and 9 in mice and each bind specific sialic acid linkages on glycans. The demonstration that adipose upregulates several STases and increases both ⍺ -2,3 and ⍺-2,6 linked sialic acids is critical because these linkages control which Siglec receptor is activated and suggests that adipose-induced sialylation can potentially lead to activation of more than one Siglec receptor consequently promoting an immunesuppressive tumor immune microenvironment.

An interesting finding is that the tumorigenic capacity of hyposialylated OC cells is curtailed by the presence of T cells. Tumors formed by TKO^SNAlow^ cells in athymic nude mice, still however, grew significantly slower compared to tumors from TKO^SNAhigh^ cells. These results demonstrate that although immune cells such as NK, B cells and macrophages, for instance, can significantly delay the growth of TKO^SNAlow^cells *in vivo*, T cells are required to fully prevent their tumorigenic capacity. Moreover, the presence or absence of T cells seems to be the switch that dictates whether tumors will form or whether hyposialylated OC cells will be reprogrammed to be hypersialylated thus lead to successful tumor formation.

In conclusion, we put forth a proposed model for adipose -induced sialic acid reprogramming (**Fig. 9**). In our model, hypersialylated OC cells that migrate to adipose-rich sites such as the omentum readily establish in this niche and upregulate STase expression and overall cell surface sialylation. In contrast, hyposialylated OC cells become targets of the immune system. In the absence of T cells however, hyposialylated OC cells undergo sialylation reprogramming, which further selects for highly sialylated OC cells as a “feed-forward” effect, which can contribute to an immunesuppressive tumor microenvironment.

**Figure 9.**
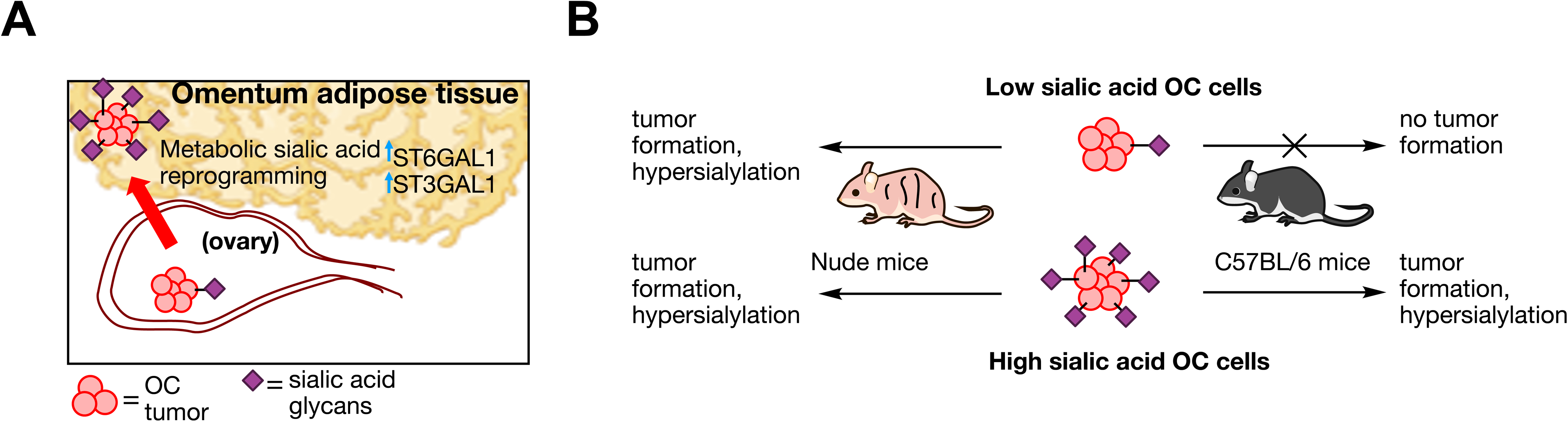
Proposed model of sialylation reprogramming by adipose microenvironment. **A.** Working model for OC sialic acid reprograming in adipose-rich niches involving upregulation of several SiaTs. **B**. OC cell sialylation dictates tumor formation in an immune-dependent manner. Absence of T cells allow tumor formation and sialylation reprogramming in hyposialylated OC cells.

## 7 Author Contributions

AF: investigation, methodology, validation, writing – review and editing

GL: investigation, methodology, validation, writing – original draft

NA: data curation, formal analysis

TW: resources, investigation

RT: investigation

SS: investigation

RG: resources

RM: resources, funding acquisition

GM: funding acquisition

CF: conceptualization, funding acquisition, supervision, writing – original draft, review and editing

AA: conceptualization, funding acquisition, supervision, project administration, writing – original draft, review and editing

## 9 Acknowledgments

The authors acknowledge The Microscopy, Imaging and Cytometry Resources Core and The Biobank and Correlative Sciences Core at Karmanos Cancer Institute (supported in part by NIH Center grant P30 CA22453 to the Karmanos Cancer Institute). The authors also wish to thank the patients and clinical staff for providing resources used in this study.

## 10 Supplementary Material

**Supplementary Figure 1:** Full blots for western blot data shown in Figure 1D.

**Supplementary Figure 2: Lectin panel for human OC lines.** Human OC cell lines (OCSC1-F2, R182, OVCAR3, OVCA432) were stained with SNA-FITC, Mal-I-FITC, Mall II-FITC, or PNA-FITC.

**Supplementary Figure 3: Lectin panel for mouse OC cell lines.** Mouse OC cell lines (TKO and ID8*Trp53*-/-) were stained with SNA-FITC, Mal-I-FITC, Mall II-FITC, or PNA-FITC.

**Supplementary Figure 4: SNA staining for different passages of TKO mouse OC cell line.** Representative SNA staining profiles on passages of TKO ovarian cancer cells cultured between December, 2022 and March, 2023.

**Supplementary Figure 5:** Results of Short tandem repeat (STR)-based authentication of TKO^SNAlow^ vs TKO^SNAhigh^.

**Supplementary Figure 6: Lectin panel of TKO^SNAhigh^ and TKO^SNAlow^.** Parental TKO mouse OC cells were sorted based on SNA levels and each subpopulation was stained with SNA-FITC, Mal-I-FITC, Mall II-FITC, or PNA-FITC.

**Supplementary Figure 7: A.** Model showing recognition site for PNA before and after neuraminidase treatment. **B.** TKO^SNAhigh^ and TKO^SNAlow^ cells were stained with PNA before after treatment with neuraminidase.

## 11 Data Availability Statement

The datasets generated from this study will be made publicly available upon acceptance of the manuscript.

