## Supplementary Materials for "Adipose microenvironment promotes hypersialylation of ovarian cancer cells"

### *Supplementary Material*

**Table of contents**

**Supplemental figures**

Full Western blot images S3

Lectin panels S4-S5

Short tandem repeat-based authentication of TKO SNA low vs TKO SNA high. S6

Sorting of TKO SNA low and TKO SNA high cell subpopulations S7

Determination of relative sialic acid levels in TKO SNA high vs low cells S8

**Supplementary Data**

#### 1 Full Western blot images:

Full western blots used in the paper. Red box denotes region that was cropped for figures.

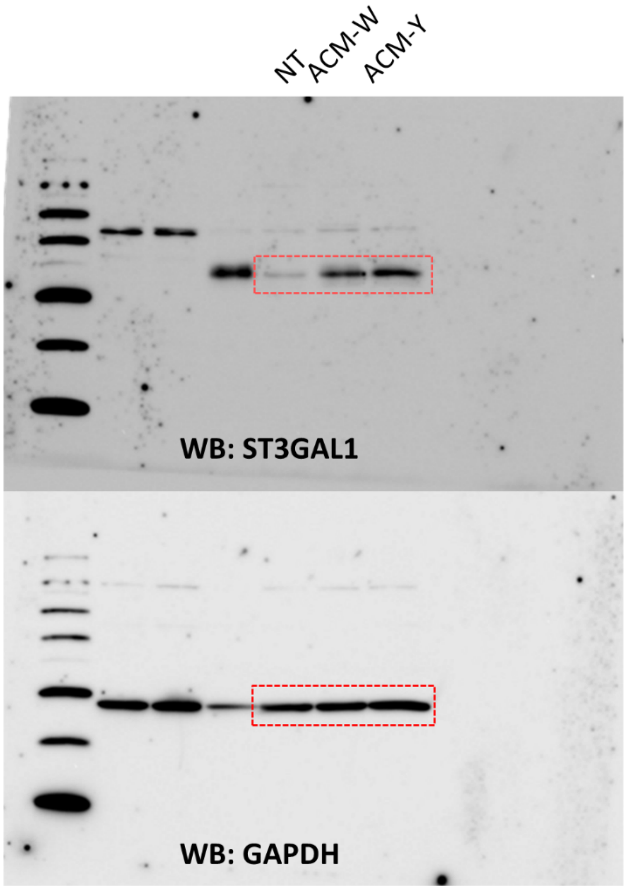

**Supplementary Figure 1:** Full blots for western blot data shown in Figure 1D.

#### 2 Lectin panel of ovarian cancer cell lines:

Normal cultured cell lines OCSC1-F2, R182, OVCAR3 and OVCA432, TKO or ID8p53KO were stained using SNA, Mal-I, Mal- II, and PNA. Cultured cells were trypsinized for 4 minutes using 0.25% trpyspin. This was then diluted using culture media before centrigation at 300 g for 5 minutes. Culture media was decanted before addition of FACs buffer to resuspend cells. Cells were then passed through a 70uM filter to reduce cell clumping. Cells when then pelleted at 300 g for 5 minutes. FACs buffer was decanted and cells were then stained with 500 µL of 1 : 400 SNA, Mal-I, Mal-II, PNA, or PBS. Cells were stained on ice for 45 minutes. After staining, 1 mL of FACs was added before pelleting and decanting liquid from pellets. This was repeated three times. 500 µL was added to resuspend cells for flow cytometry analysis.

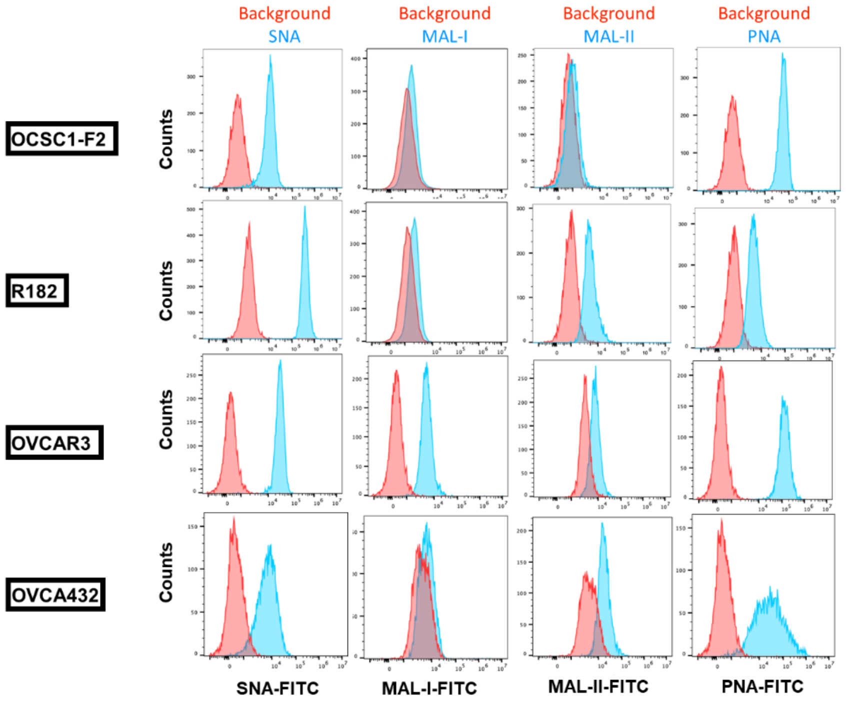

**Supplementary Figure 2: Lectin panel for human OC lines.** Human OC cell lines (OCSC1-F2, R182, OVCAR3, OVCA432) were stained with SNA-FITC, Mal-I-FITC, Mall II-FITC, or PNA-FITC.

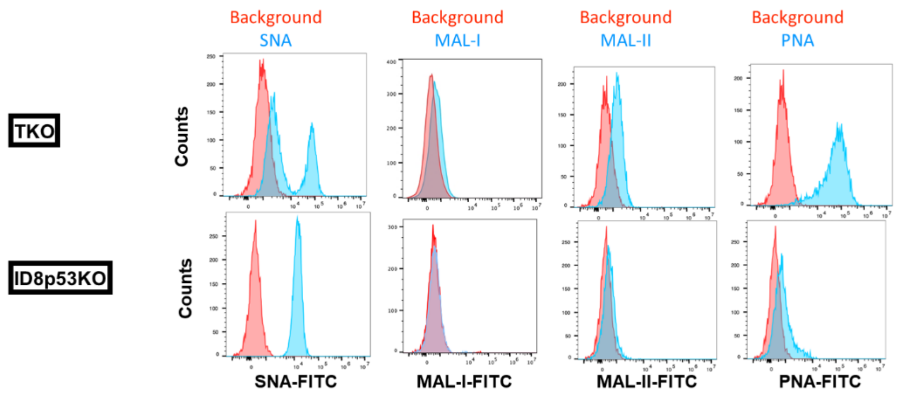

**Supplementary Figure 3: Lectin panel** for **mouse OC cell lines.** Mouse OC cell lines (TKO and ID8*Trp53*-/-) were stained with SNA-FITC, Mal-I-FITC, Mall II-FITC, or PNA-FITC.

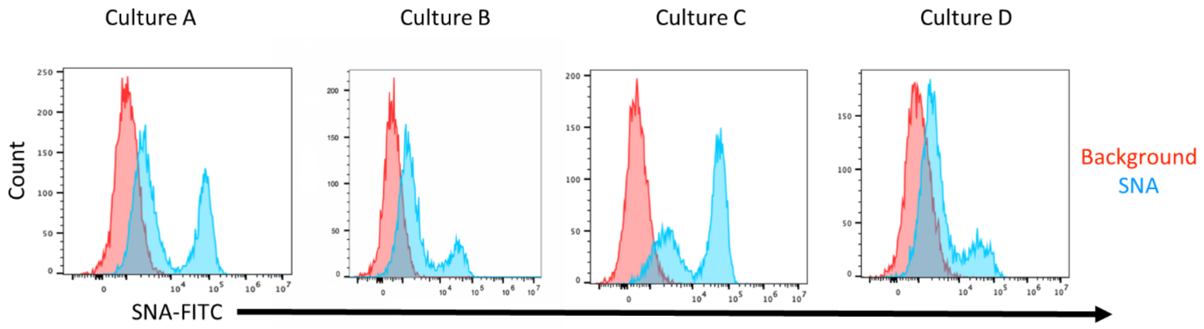

**Supplementary Figure 4: SNA staining for different passages of TKO mouse OC cell line.** Representative SNA staining profiles on passages of TKO ovarian cancer cells cultured between December, 2022 and March, 2023.

MCLAMC.R2.0423

labcorp

Mouse Cell Line Authentication Match Calculator

Please note: Homozygous alleles count as ONE; only input alleles that are present in <u>both</u> the Questioned and Reference Samples; Samples with more than (5) alleles in any one marker is an indication of a mixture and/or severe genetic instability and may not be suitable for authentication calculations (discard stock and restart/purchase new stock or consult your cell line testing representative for more help)

| | QUESTIONED SAMPLE | | | | | REFERENCE SAMPLE | |
|---|---|---|---|---|---|---|---|
| Cell Line Name | TKO sna high | | | | | TKO sna low | |
| Source | p10.1 | | | | | p10.1 | |
| Testing Date | 7/1/2023 | | | | | 7/1/2023 | |
| MARKER | ALLELES | | | | | ALLELES | |
| M18-3 | 16 | | | | | 16 | 17 |
| M4-2 | 20.3 | | | | | 20.3 | |
| M6-7 | 15 | 16 | | | | 15 | |
| M19-2 | 13 | | | | | 13 | |
| M1-2 | 19 | | | | | 19 | |
| M7-1 | 25.2 | | | | | 25.2 | |
| M1-1 | 14 | | | | | 14 | |
| M3-2 | 14 | | | | | 14 | |
| M8-1 | 16 | | | | | 16 | |
| M2-1 | 9 | 16 | | | | 9 | 16 |
| M15-3 | 22.3 | 23.3 | | | | 22.3 | 23.3 |
| M6-4 | 16 | 18 | | | | 16 | 18 |
| M11-2 | 16 | | | | | 16 | |
| M17-2 | 12 | | | | | 12 | |
| M12-1 | 17 | | | | | 17 | |
| M5-5 | 14 | 15 | | | | 14 | |
| MX-1 | 27 | | | | | 27 | |
| M13-1 | 17 | | | | | 17 | |

| | |
|---|---|
| Percent Match (ATCC SDO ASN-0002-2022; Tanabe et al) | 93.33% |
| Masters Algorithm | 91.30% |

Percent Match (ASN-0002-2022; Tanabe et al) =2(allele matches)/(number question alleles +number reference alleles)

Masters Algorithm = (allele matches)/(number question alleles)

AUTHENTICATION STATUS

A "Percent Match":

- **100% = <u>Authenticated</u>
- **Between 80%-99%** = Result is consistent with two samples being related
- **Between 56-79%** = Result is inconclusive and may need further testing using alternative methods
- **Between 0-55%** = Result is consistent with the two samples being unrelated

**NOTE: At this time, there is NO universally accepted mouse cell line authentication standard and thus there is no currently accepted calculation for true mouse cell line authentication. NIST is currently in the process of establishing a calculation that takes mouse in-breeding into consideration, without that consideration, two mouse cell lines derived from the same STRAIN of mouse could look very similar in their STR profile results. Following months of rigorous testing by our lab and with the help of clients submitting mouse cell lines for STR profiling, we recommend that mouse cell lines be "authenticated" only when there is an exact match (100%) to the known reference STR profile – if you have questions on this, please contact your Labcorp representative.

***References***

1. *ANSI/ATCC ASN-0002-2022 REVISION. Authentication of Human Cell Lines: Standardization of STR Profiling.*
2. *Tanabe H et al. Cell line individualization by STR multiplex system in the cell bank found cross-contamination between ECV304 and EJ-1/T24. Tiss Cult Res Commun 1999;18:329-38.*
3. *Masters JR et al. Short tandem repeat profiling provides an international... Proc Natl Acad Sci USA 98(14):8012-8017, 2001*

Labcorp • 1440 York Court, Burlington, NC 27215 USA

• www.celllineauthentication.com •

1-800-IDENTITY® (433-6848) [USA] • 001-866-FAMILIA (326-4542) [Mexico] • +1-513-985-9777 [International]

**Supplementary Figure 5:** Results of Short tandem repeat (STR)-based authentication of TKO$^{SNAlow}$ vs TKO$^{SNAhigh}$.

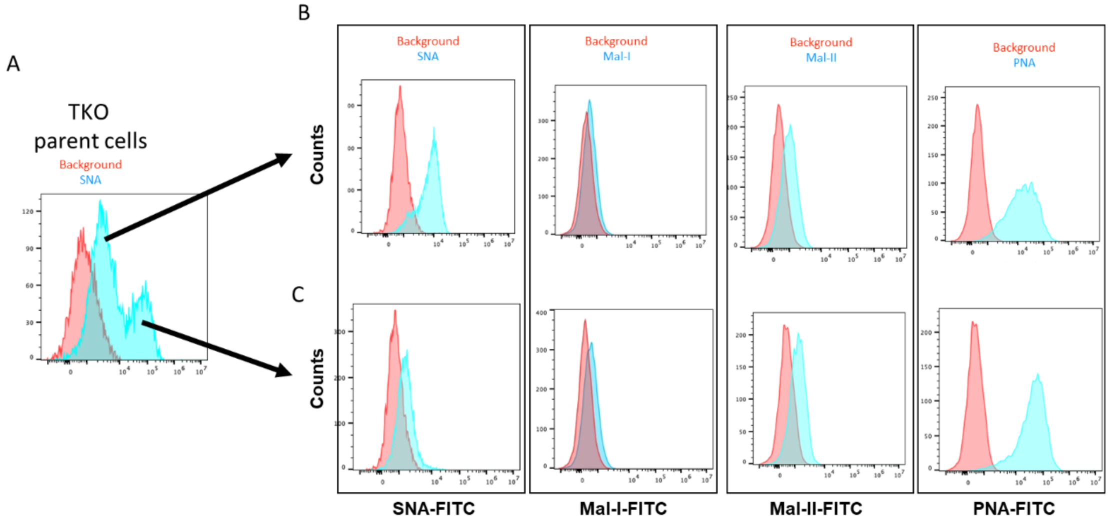

**Supplementary Figure 6: Lectin panel of TKO$^{SNAhigh}$ and TKO$^{SNAlow}$.** Parental TKO mouse OC cells were sorted based on SNA levels and each subpopulation was stained with SNA-FITC, Mal-I-FITC, Mall II-FITC, or PNA-FITC.

**Determination of relative sialic acid levels in TKO high vs TKO low via neuraminidase treatment.**

To determine the relative difference in overall sialic acid between TKO high and TKO low, these ovarian cancer cell lines were subjected to neuraminidase treatment. Cells were grown in T75 culture flask in DMEM/F12 supplemented with l-Glutamine, 10% FBS, and 1% penstrep. Cells were trypsinized using 0.25% trypsin for 3 minutes and this was removed via media dilution and removal of the supernatant after centrifugation. Cells were counted and 1x10^6 cells were transferred to 15 mL falcon tubes. Cells were treated with 20 units of neuraminidase A (new England biolabs P0722S) in 1 mL of culture media. A separate control was also generated containing no neuraminidase. Both of these tubes were allowed to rotate at 37 °C for 2 hrs. After this treatment cells were rinsed 3 times with PBS containing 10% FBS and 1% pen/strep. Cells were then subjected to PNA-FITC staining at a 1:400 dilution for 45 minutes on ice. Cells were then rinsed 3 times with PBS before flow cytometry.

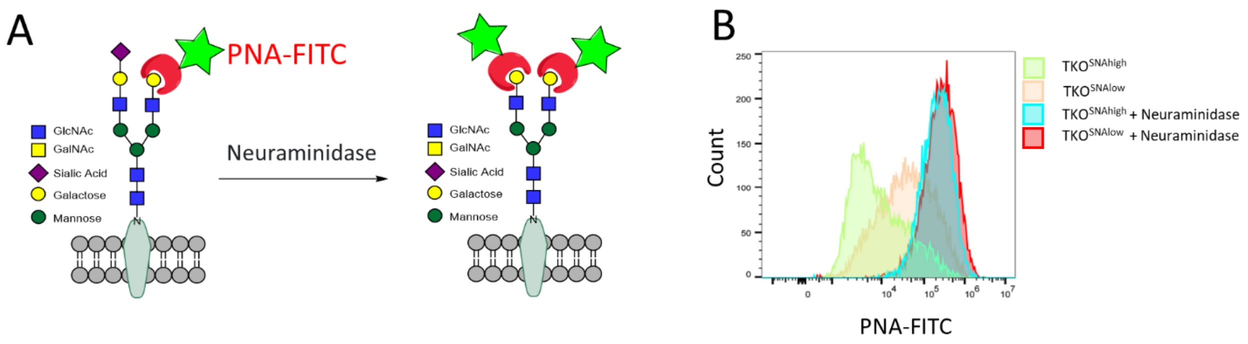

**Supplementary Figure 7: A.** Model showing recognition site for PNA before and after neuraminidase treatment. **B.** TKO$^{SNAhigh}$ and TKO$^{SNAlow}$ cells were stained with PNA before after treatment with neuraminidase.
